# Algal photosynthesis converts nitric oxide into nitrous oxide

**DOI:** 10.1101/745463

**Authors:** Adrien Burlacot, Pierre Richaud, Arthur Gosset, Yonghua Li-Beisson, Gilles Peltier

## Abstract

Nitrous oxide (N_2_O), a potent greenhouse gas in the atmosphere, is produced mostly from aquatic ecosystems, to which algae substantially contribute. However, mechanisms of N_2_O production by photosynthetic organisms are poorly described. Here, we show that the green microalga *Chlamydomonas reinhardtii* reduces NO into N_2_O using the photosynthetic electron transport. Through the study of *C. reinhardtii* mutants deficient in flavodiiron proteins (FLVs) or in a cytochrome p450 (CYP55), we show that FLVs contribute to NO reduction in the light, while CYP55 operates in the dark. Furthermore, NO reduction by both pathways is restricted to Chlorophytes, organisms particularly abundant in ocean N_2_O-producing hotspots. Our results provide a mechanistic understanding of N_2_O production in eukaryotic phototrophs and represent an important step toward a comprehensive assessment of greenhouse gas emission by aquatic ecosystems.

**One sentence summary:** Green microalgae produce N_2_O using flavodiiron proteins in the light and a cytochrome P450 NO reductase in the dark.

## INTRODUCTION

Although nitrous oxide (N_2_O) is present in the atmosphere at concentrations 1,000 times lower than CO_2_, it is recognized as the third most potent greenhouse gas after CO_2_ and CH_4_, contributing to ∼6% of the total radiative forcing on earth (*1, 2*). In addition, N_2_O is the main ozone-depleting gas produced in our planet (*3*). Since 1970, N_2_O concentration in the atmosphere has been rising, reaching the highest measured production rate in the past 22,000 years (*2*). Natural sources of N_2_O account for 64% of the global N_2_O production, mostly originating from soils and oceans (*4*). Bacteria and fungi widely contribute to this production, N_2_O being produced during nitrification by the reduction of nitric oxide (NO) mediated by NO reductases (NORs) (*5, 6*).

Microalgae are primary biomass producers in oceans and lakes. For decades, N_2_O production has been detected in samples from the ocean, lakes and coastal waters but the respective contribution of prokaryotic and eukaryotic organisms to this phenomenon remains to be investigated (*7-9*), and particularly the contribution of microalgae has been overlooked (*7, 10*). Algal blooms correlate with N_2_O production (*8, 10, 11*), and recently, axenic microalgal cultures were shown to produce substantial amount of N_2_O (*12-14*). Despite the possible ecological significance of microalgae in N_2_O emissions (*10*), little is known about the molecular mechanisms of N_2_O production in microalgae.

In bacteria, membrane-bound NORs belong to the haem/copper cytochrome oxidase family (*15*), some of which containing a c-type cytochrome (*16*). Soluble bacterial NORs belong to the flavodiiron family (FLVs), enzymes able to reduce NO and/or O_2_ (*17*). In fungi, NORs are soluble and belong to the cytochrome P450 (CYP55) family (*18, 19*). The genome of the model green microalga *Chlamydomonas reinhardtii* harbors both a CYP55 fungal homolog (*12*) and FLVs bacterial homologs (*20*). Based on RNAi silencing experiments in *C. reinhardtii*, Plouviez *et al*. concluded that the CYP55 homolog is the main contributor to N_2_O production (*13*). However, this gene has so far only been identified in three sequenced algal genomes (*21*) suggesting the occurrence of other mechanisms. *C. reinhardtii* FLVs have been recently shown to be involved in O_2_ photoreduction (*22*), but have not been considered as catalyzing NO reduction so far (*23*). Thus, the major players involved in N_2_O production in microalgae remain to be elucidated.

In this work, by measuring NO and N_2_O gas exchange using a Membrane Inlet Mass Spectrometer (MIMS) during dark to light transitions, we report on the occurrence of a PSI-dependent photoreduction of NO into N_2_O in the unicellular green alga *C. reinhardtii.* Through the study of mutants deficient in FLVs or CYP55 or both, we conclude that FLVs mainly contributes to N_2_O production in the light, while CYP55 is mostly involved in the dark. The ecological implication of NO reduction to N_2_O by microalgae, a phenomenon shown to be restricted to algae of the green lineage, is discussed.

## RESULTS

### *C. reinhardtii* reduce NO to N_2_O in the light using the photosynthetic electron transport chain

Measurements of NO and N_2_O exchange were performed on *C. reinhardtii* cell suspensions using MIMS. To ensure sufficient amount of substrate, we injected exogenous NO into the cell suspension. Because NO can be spontaneously oxidized into nitrite in the presence of O_2_, experiments were performed under anoxic conditions by adding glucose and glucose oxidase to the cell suspension as an O_2_ scavenger. After NO injection, NO uptake and N_2_O production were measured in the dark (**Fig. 1A and B**), and the NO uptake rate was found about twice higher than the N_2_O production rate (**Fig. 1C**), thus indicating the existence in the dark of a stoichiometric reduction of NO into N_2_O. Upon illumination, a strong increase in both NO uptake and N_2_O production was observed (**Fig. 1A and B**). The NO_uptake_/N_2_O_production_ ratio increased up to a value of 2.5 (**Fig. 1C**). This likely indicates the occurrence of two distinct phenomena in the light: *i*. a stoichiometric photoreduction of NO into N_2_O, and *ii*. a photo-dependent NO uptake process independent of N_2_O production.

**Figure 1.**
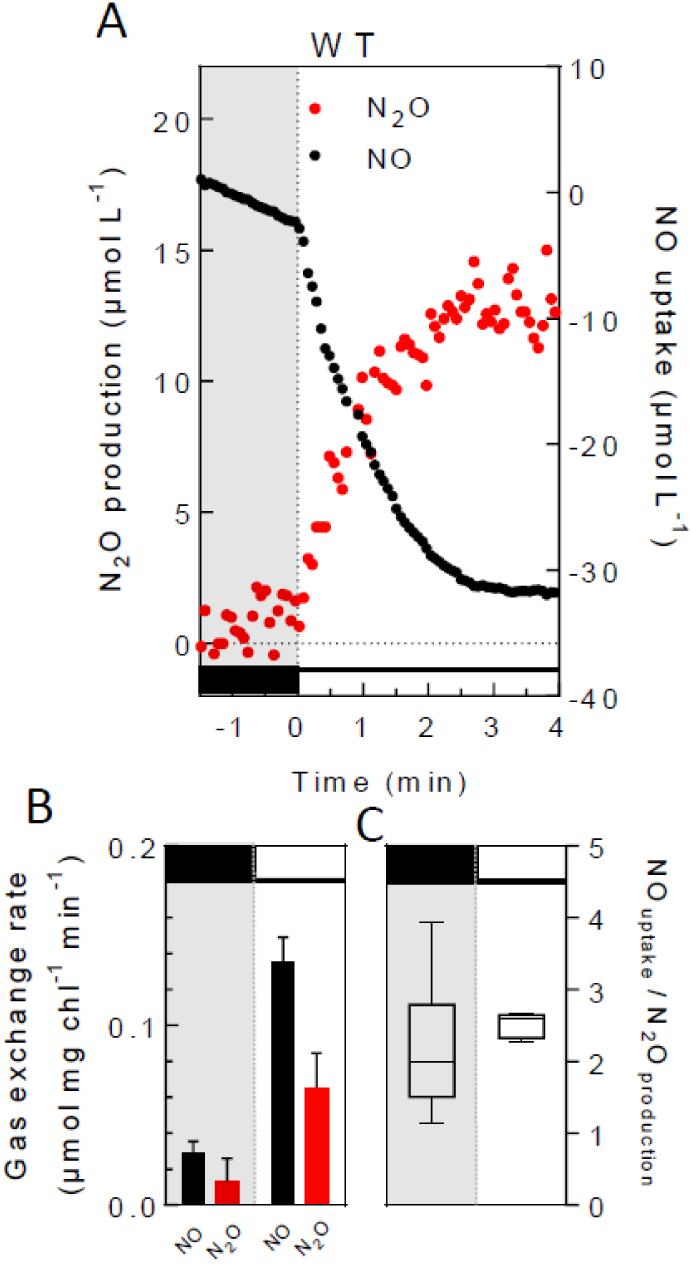
Reduction of NO into N_2_O in the green alga *C. reinhardtii*. After 1 min anaerobic acclimation, NO was injected in the cell suspension to a final concentration of 45 µM. After 3 min in the dark, cells were illuminated with green light (3,000 µmol photon m^−2^ s^−1^). (**A**) Representative traces of cumulated amounts of NO uptake (black circles) and N_2_O production (red circles) measured in the control *C. reinhardtii* strain during a dark to light transient. (**B**) Dark (left panel) and light-dependent (right panel) NO uptake rates (black) and N_2_O production rates (red). Data shown are mean values ± SD (n=4). (**C**) Box plot of the ratio of NO uptake rate over N_2_O production rate in the dark and over the entire light period. (mean, min, max, n=8).

In order to determine whether photosynthesis could serve as a source of electrons for NO reduction in the light, we studied the effect of two inhibitors, 3,4-dichlorophenyl-1,1-dimethylurea (DCMU), a potent photosystem II (PSII) inhibitor, and 2,5-dibromo-3-methyl-6-isopopyl-*p*-benzoquinone (DBMIB), a plastoquinone analog blocking the photosynthetic electron flow between PSII and PSI. While both inhibitors had no effect in the dark, DCMU decreased the light-dependent NO uptake and the light-dependent N_2_O production rate by 65% and 55%, respectively (**Fig. 2A, C, D**). On the other hand, DBMIB decreased the light-dependent NO uptake rate by more than 90% and completely abolished the light-dependent N_2_O production (**Fig. 2B, C, D**). The above experiments performed with photosynthetic inhibitors point to the involvement of PSI in the photoreduction of NO into N_2_O, the partial inhibition observed in the presence of DCMU likely resulting from the occurrence of a non-photochemical reduction of plastoquinones by the alternative NAD(P)H dehydrogenase 2 (NDA2), as previously reported for *C. reinhardtii* (*24*). We conclude from these experiments that *C. reinhardtii* can reduce NO into N_2_O in a light-dependent manner using electrons provided by the photosynthetic electron transport chain.

**Figure 2.**
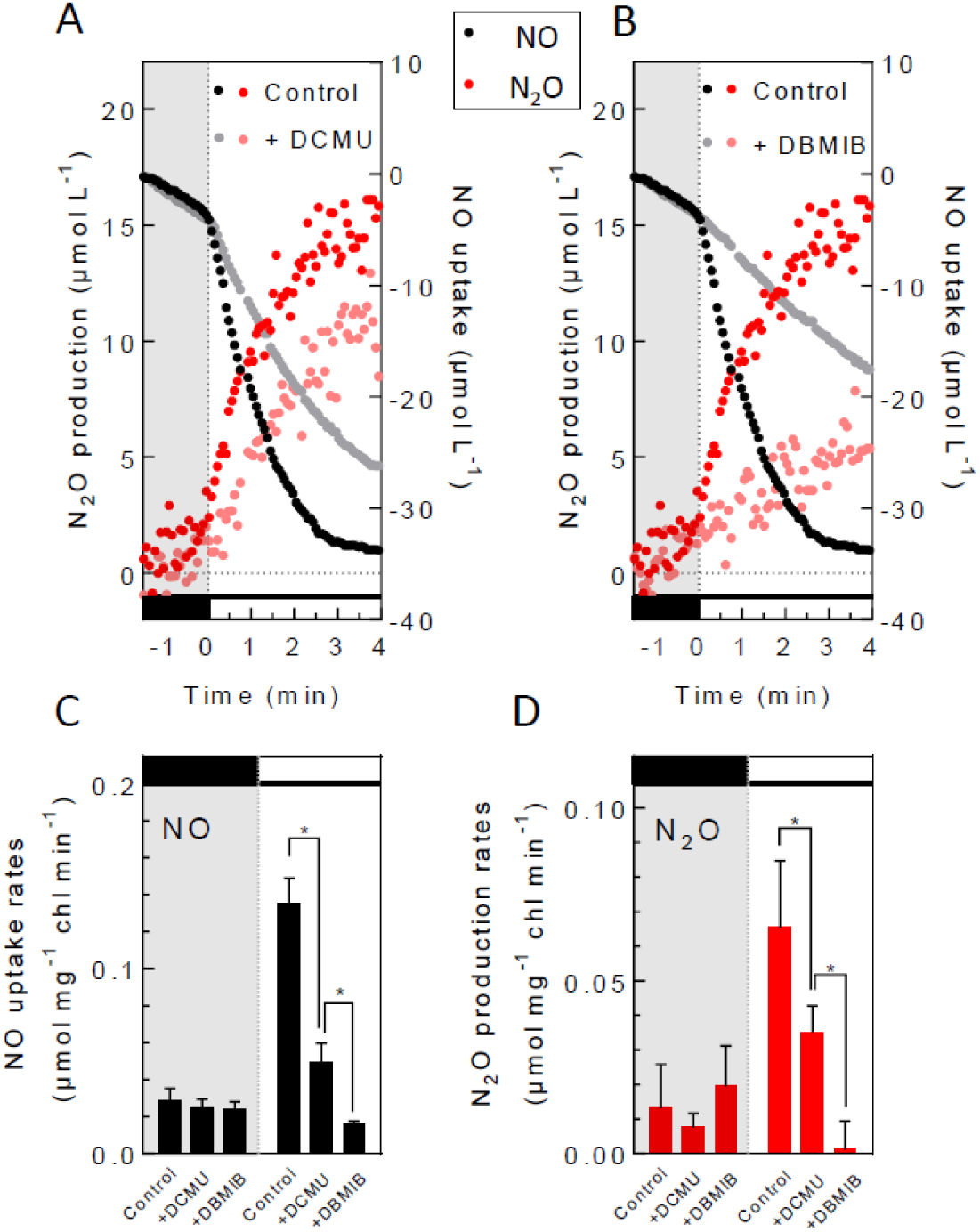
The photoreduction of NO into N_2_O involves the photosynthetic electron transport chain. NO and N_2_O gas exchange were measured during a light transient as described in **Fig. 1** in the absence or presence of photosynthesis inhibitors DCMU (10 µM) or DBMIB (2 µM). (**A, B**) Representative traces of cumulated NO uptake (black dots) and N_2_O production (red dots) in the control strain in the absence (full color dots) or presence of DCMU or DBMIB (greyed out dots). (**C**) Dark (left panel) and light-dependent (right panel) NO uptake rates measured in the absence or presence of inhibitors. (**D**) Dark (left panel) and light-dependent (right panel) N_2_O production rates measured in the absence or presence of inhibitors. Data shown are mean values ± SD (n=4). Asterisks mark significant differences (*P* < 0.05) based on multiple T-tests.

### Light-dependent N_2_O photoproduction mostly relies on FLVs

Depending on their origin, bacterial flavodiiron proteins are capable of catalyzing O_2_ reduction, NO reduction or both reactions (*23*). In *C. reinhardtii*, two FLVs (FLVA and FLVB) have recently been described as catalyzing O_2_ photo-reduction using the reducing power produced at the PSI acceptor side by photosynthesis (*22*). In order to determine the contribution of FLVs to NO photoreduction, we analyzed NO and N_2_O exchange during dark to light transients in three previously characterized *flvB* mutants (*22*). These mutants are impaired in the accumulation of both FLVB and FLVA subunits (*22, 25, 26*). In the dark, no significant difference in NO reduction neither in N_2_O production were found between the three *flvB* mutant as compared to the control (**Fig. 3A, C, E, F**; **Supplemental Fig. S1A and B**). In contrast, the N_2_O production and NO uptake induced by light were respectively decreased by 70% and 50% in *flvB-21* as compared to the control strain (**Fig. 3A, C, E, and F**). Similar effects were observed in the two others independent *flvB* mutants (**Supplemental Fig. S1A and B**). We conclude from this experiment that FLVs are involved in the light-dependent reduction of NO into N_2_O, using electrons produced by photosynthesis.

**Figure 3.**
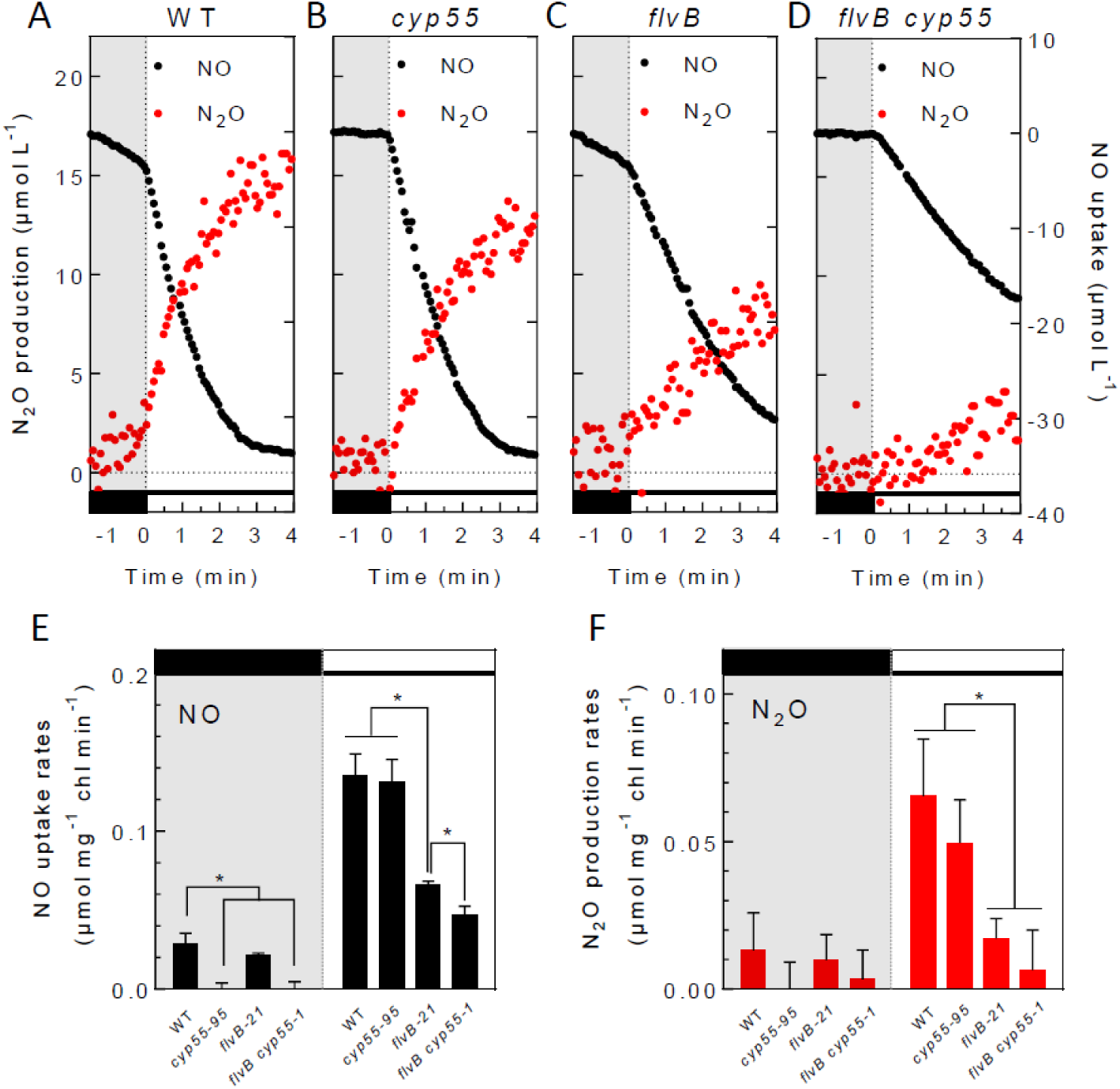
The light-dependent N_2_O production involves FLVs while the dark production involves CYP55. NO and N_2_O gas exchange were measured during a light transient as described in **Fig. 1** in the control strain (**A**), the *cyp55-95* mutant deficient in CYP55 (**B**), in the *flvB-21* mutant deficient in FLVB (**C**), and in a double *flvB cyp55-1* mutant (**D**). (**A-D**) Representative traces of cumulated NO production (black) and N_2_O production (red). (**E**) Dark (left panel) and light-dependent (right panel) NO uptake rates. (**F**) Dark (left panel) and light-dependent (right panel) N_2_O production rates. Data shown are mean values ± SD (n=4). Asterisks mark significant differences (*P* < 0.05) based on multiple T-tests.

### Dark N_2_O production relies on CYP55

A homolog of the nitric oxide reductase from *Fusarium oxysporum* (CYP55) encoded by the *C. reinhardtii* genome (Cre01.g007950) was recently proposed to be involved in NO reduction (*13*). In order to investigate the contribution of the CYP55 homolog to N_2_O production, we obtained three *C. reinhardtii* insertion mutants from the CLiP library (https://www.chlamylibrary.org; (*27*). Two of these putative *cyp55* mutants have been predicted to hold an insertion of the paromomycin resistance cassette in introns, while the third one has a predicted insertion in an exon (*cyp55-95*) (**Supplemental Table S1; Supplemental Fig. S2A**). Positions of insertions were confirmed by PCR on genomic DNA (**Supplemental Fig. S2B and C**). Both NO uptake and N_2_O production in the dark were completely abolished in all three *cyp55 mutants* (**Fig. 3 A, B, E, F**; **Supplemental Fig. S1**). However, rates of NO reduction and N_2_O production induced by light were not significantly affected in all three *cyp55* mutants (**Fig. 3 A, B, E, F; Supplemental Fig. S2**). We conclude from this experiment that CYP55 is responsible for the entire reduction of NO to N_2_O in the dark.

Furthermore, a double mutant with mutations in both *CYP55* and *FLVB* was obtained by crossing *cyp55-95* and *flvB-21* strains, and three independent progenies (*flvB cyp55-1, -2* and *-3*) were isolated (**Supplemental Fig. S3**). The N_2_O production was nearly abolished in these double mutant lines both in the dark and in the light (**Fig. 3D, F; Supplemental Fig. S4B**). The quantity of N_2_O produced during the first minute of illumination was significantly lower in the double mutants when compared to the single *flvB-21* mutant (**Supplemental Fig. S5**) so was the NO uptake rate (**Fig. 3E; Supplemental Fig. S4A**). Thus, in the absence of FLVs, CYP55 contributes to the light-dependent NO reduction to N_2_O production.

### N_2_O production is restricted to Chlorophytes and correlates with the presence of *FLV* and *CYP55*

We then explored the ability of microalgae originating from different phyla to reduce NO to N_2_O either in the dark or in the light. We focused on species with sequenced genomes or transcriptomes and listed the presence of *FLV* and *CYP55* homologous genes in the different species (**Supplemental Table S3**). The ability to produce N_2_O in the dark was only found in algae of the green lineage (Chlorophytes) (**Supplemental Fig. S6**) and associated to the presence of a *CYP55* gene homolog (**Fig. 4**). A light-dependent N_2_O production was also only found in Chlorophytes (**Supplemental Fig. S6**) and associated to the presence of *FLV* genes (**Fig. 4**). Note that diatoms and red algae showed a light-dependent NO uptake but without significant N_2_O production (**Supplemental Fig. S6E-I**). Similar light-dependent NO uptake was also evidenced in the *C. reinhardtii flvB cyp55* double mutants (**Fig. 3; Supplemental Fig. S4**), thereby highlighting the existence in all analyzed algal species of a mechanism of photo-dependent NO uptake which likely reflects oxidation of NO by the PSII-produced O_2_. We conclude from these experiments that N_2_O production in algae mostly relies on CYP55 in the dark and on FLVs in the light.

**Figure 4.**
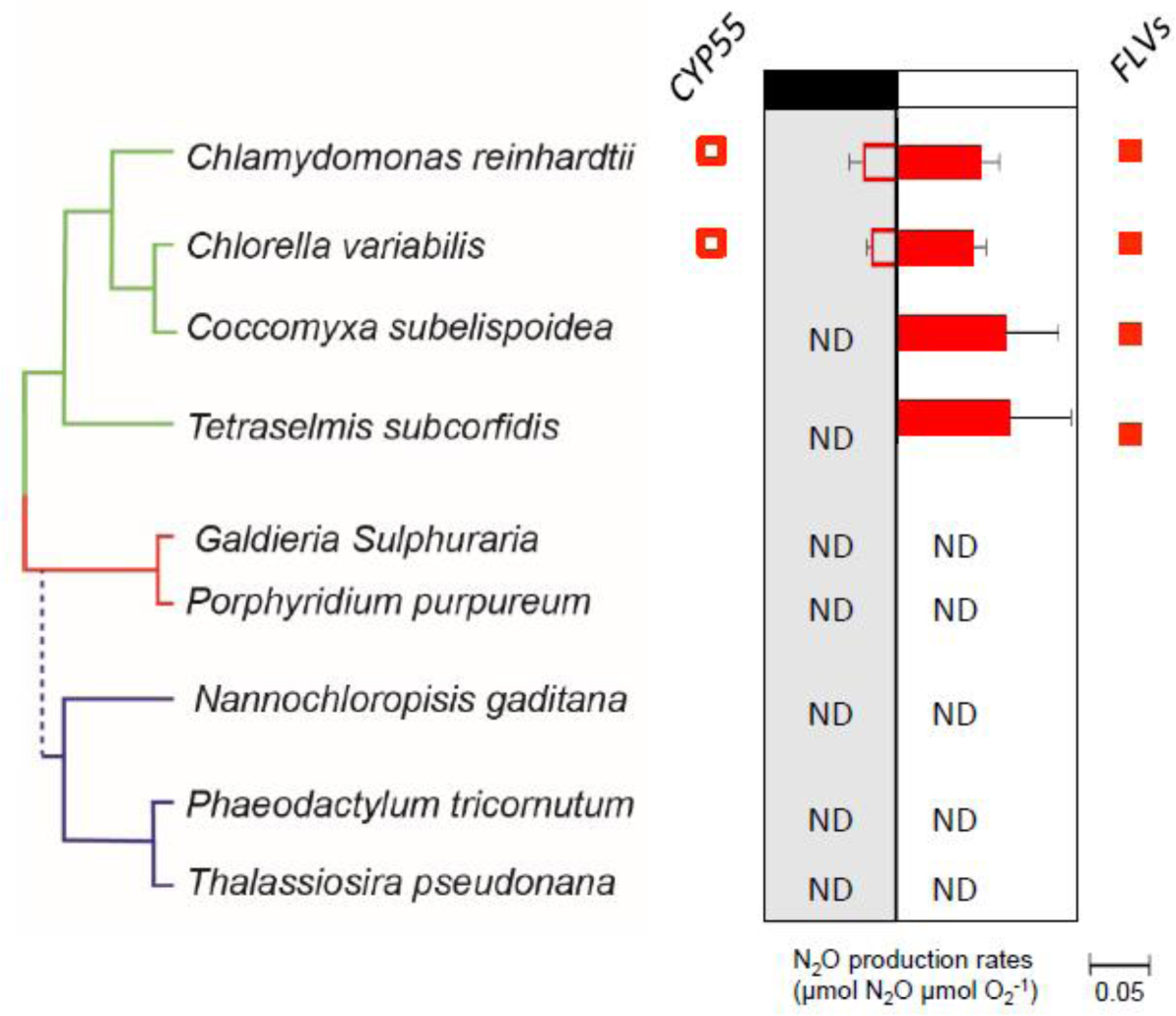
The light-dependent N_2_O production is associated to the presence of *FLVs* while N_2_O production in the dark is associated to the presence of *CYP55* in algal genomes. Presence of *CYP55* and *FLV* homologous genes in different algal species are respectively shown by open or filled red squares. NO and N_2_O gas exchange were measured in the different species during a dark-light transient as described in **Fig. 1**. Maximal gross O_2_ production rate by PSII were measured on the same samples using labeled ^18^O_2_. When detected, N_2_O production rates were normalized to the corresponding maximum gross O_2_ production rates. Relative rates of dark (left panel) and light-dependent (right panel) N_2_O production are shown. Data shown are mean values ± SD (n=3).

## DISCUSSION

Although the ability of green algae to produce N_2_O has been documented for more than 30 years (*28*), molecular mechanisms remained enigmatic. In this work, we show that the green microalga *C. reinhardtii* can reduce NO to N_2_O in the dark as well as in the light but employing different mechanisms. The light-dependent reduction of NO is catalyzed by FLVs and uses electrons produced by the photosynthetic chain while the dark reaction is mediated by CYP55. We have further shown that the ability to produce N_2_O is restricted to Chlorophytes, in which dark and light-dependent productions respectively correlate with the presence of *CYP55* and *FLV* genes.

In microalgae and land plants, the photosynthetic electron flow is principally used to reduce CO_2_ via the Calvin-Benson cycle. Several alternative electron fluxes also occur, which usually play critical roles during acclimation of photosynthesis to environmental changes (*29, 30, 31*). Among known electron acceptors of alternative electron fluxes, one finds *i*. molecular O_2_ that can be reduced into water or reactive oxygen species by different mechanisms (*32*), *ii*. protons that can be reduced into dihydrogen by hydrogenases (*33*), and *iii*. nitrites reduced into ammonium by nitrite reductases (*34*). We provide evidence here that NO is to be considered as a new electron acceptor of photosynthesis downstream PSI. The maximal proportion of the photosynthetic electron flow used for NO reduction is estimated around 5% considering that at PSII four electrons are produced per molecule of O_2_ released, and that two electrons are used per molecule of N_2_O produced. Due to this relatively low rate, it seems unlikely that NO photoreduction significantly functions as a valve for electrons during the functioning of photosynthesis, as it has been shown for O_2_ photoreduction by FLVs under aerobic conditions (*22*) or H_2_ photoproduction under anaerobic conditions (*35*). Nevertheless, NO photoreduction could play a regulatory role during anaerobic photosynthesis, as shown for other minor pathways such as chlororespiration under aerobic conditions (*30, 36*). In photosynthetic organisms, FLVs have so far been solely involved in oxygen reduction, this electron valve being critical for growth under fluctuating light conditions (*22, 26, 37, 38*). We show here that FLVs play a dual function in the reduction of O_2_ as well as NO in chlorophytes, and that the significance of this duality on the functioning and regulation of photosynthesis should be considered in future studies.

In microalgae, N_2_O production occurs mostly during nitrogen assimilation, when nitrates or nitrites are supplied as nitrogen sources (*12*). During this process, nitrite is reduced into ammonium by the nitrite reductase (*34*), but can alternatively be reduced to NO by either a NO-forming nitrite reductase (*39*) or by respiratory oxidases (*40*). N_2_O was observed in conditions where NO is produced and CYP55 was suggested to be involved in its reduction (*13*). Our work establishes that the CYP55 is indeed involved in this process, but essentially in the dark. Production of N_2_O by microalgal cultures was also reported to greatly vary depending on culture conditions like illumination, nitrogen source or oxygen availability (*10, 12, 28*). Although this variability could have been due to differences in the strains used or fine regulatory mechanisms, it remained unexplained (*13*). Our results clearly demonstrate the existence of two distinct pathways of NO reduction. Thus, variability in N_2_O production may result from variation in the relative importance of both CYP55 and FLVs reductive pathways depending on experimental conditions.

NO is a known signal molecule involved in various regulatory mechanisms in all living organisms. In microalgae, NO is involved in the regulation of the nitrate and nitrite assimilation pathways (*41, 42*), in the down-regulation of photosynthesis upon sulfur starvation (*43*) or in hypoxic growth (*44*). NO homeostasis results from an equilibrium between NO production by different enzymatic systems, which have been well-documented in plants (*45*), and its active degradation mediated by truncated hemoglobin that catalyze NO oxidation to nitrate in aerobic conditions (*46, 47*). Our results suggest that FLVs and CYP55, by removing NO through a reductive pathway, might be key enzymes in the control of NO homeostasis in anaerobic conditions.

Microalgae are often considered promising organisms for the production of next-generation biofuels provided that their large-scale cultivation has positive effects on the environment. However, it has recently been estimated that N_2_O produced during large-scale algal cultivation could compromise the expected environmental benefits of algal biofuels (*14*). In this context, knowledge of the molecular mechanisms involved and the selection of microalgae species with limited N_2_O production capacity are essential to limit the global warming potential of algal biofuels.

To date, the contribution of microalgae to the N_2_O atmospheric budget and global warming is not taken into account (*10*) due to our lack of knowledge about algal species concerned and the conditions of N_2_O production. We have shown that the capacity to reduce NO into N_2_O greatly depends on algal species, and is essentially restricted to Chlorophytes, the second most represented photosynthetic organisms in the ocean (*48, 49*). Chlorophytes are particularly abundant in coastal waters (*48*) where anthropic releases favor hypoxia (*50*), thus promoting N_2_O production (*51*). Coastal waters are frequently the scene of N_2_O producing hotspots following phytoplanktonic biomass accumulation (*52*) which are likely due to the existence in Chlorophytes of NO-reductive pathways involving both FLVs and CYP55. The incoming worldwide N_2_O observation network (*53*) together with follow up of phytoplanktonic communities will be decisive to better assess the microalgal contribution, particularly Chlorophytes, to the global N_2_O budget.

## Accession numbers

Genes studied in this article can be found on https://phytozome.jgi.doe.gov/ under the loci Cre12. g531900 (FLVA), Cre16.g691800 (FLVB) and Cre01.g007950 (CYP55).

Sequence data from this article can be found in the GenBank data library (https://www.ncbi.nlm.nih.gov/genbank/) under accession numbers XM_001699293.1 (FLVA), XM_001692864.1 (FLVB) and XP_001700272.1 (CYP55).

## List of abbreviations

NO: Nitrous oxide
N_2_O: Nitric oxide
FLV: Flavodiiron
PSI: Photosystem I
PSII: Photosystem II
CYP55: cytochrome p450
NDA2: NAD(P)H dehydrogenase 2

## Acknowledgments

This work was supported by the ERA-SynBio project Sun2Chem, and by the A*MIDEX (ANR-11-IDEX-0001-02) project. The authors thank Dr. Olivier Vallon for stimulating discussions and Dr. Brigitte Gontero for kindly providing the *Thalassiossira pseudonana* strain. Adrien Burlacot is a recipient of a CEA (Irtelis) international PhD studentship. The authors acknowledge the European Union Regional Developing Fund, the Region Provence Alpes Cote d’Azur, the French Ministry of Research, and the CEA for funding the HelioBiotec platform.

## Author contributions

Author contributions. A.B. and G.P. designed the experiments. A.B., P.R., and A.G. performed the experiments. A.B., Y.L.B. and G.P. analyzed the data and wrote the manuscript.

## Data and materials availability

All data needed to evaluate the conclusions in the paper are present in the paper and/or the Supplementary Materials. Additional data related to this paper may be requested from the authors.

## List of Supplementary materials

**Supplemental Table S1** *cyp55* mutant strains from the Chlamydomonas Library Project (CLiP).

**Supplemental Table S2**: Primers used to characterize *cyp55* mutants.

**Supplemental Table S3**: Eukaryotic algae used in this study and their respective genotype.

**Supplemental Figure S1**: Characterization of *cyp55* mutants

**Supplemental Figure S2**: Photoreduction of NO into N_2_O in two additional *flvB* and *cyp55* mutants.

**Supplemental Figure S3**: Characterization of *flvB cyp55* double mutants.

**Supplemental Figure S4**: Photoreduction of NO into N_2_O in *flvB cyp55* double mutants and their parental strains.

**Supplemental Figure S5**: Integrated N_2_O amounts produced during a light transient.

**Supplemental Figure S6**: NO uptake and N_2_O production in various microalgal species during a dark to light transient.

