## Supplemental materials for "Algal photosynthesis converts nitric oxide into nitrous oxide"

**Supplementary Materials:**

### MATERIAL AND METHODS

***C. reinhardtii* strains, cultivation and genetic crosses.** The *C. reinhardtii* wild-type strain CC-4533 and *flvB* mutants (*fvlB-21, fvlB-208* and *flvB-308*) were previously described (*22*). The c*yp55* mutants were obtained from the CliP library (*27*), **Supplemental Table S1** summarizes the names and predicted insertion sites in the different mutant lines. Cells were grown mixotrophically in flasks at 25°C in Tris-Acetate-Phosphate (TAP, ammonium as a nitrogen source) medium (pH 7.2) under dim light (5-10 µmol photons $m^{-2}$ $s^{-1}$). Cells were harvested during the exponential phase. Experiments presented throughout this manuscript were performed on three independent single colony-derived lines for each strain thus SDs account for standard deviation of biological triplicates.

The mutant strain $mt^-$ *flvB-21* was back-crossed with $mt^+$ CC5155 (https://www.chlamycollection.org/product/cc-5155-cw15-mt-jonikas-cmj030-f5-backcross-strain-isolate-e8/), which is a mt+ wild type strain close to the background strain used in the CliP library. The $mt^+$ *flvB-21* mutant obtained was then crossed with the $mt^-$ *cyp55-95* mutant. The progenies of this crossing were selected based on chlorophyll fluorescence to screen for the insertion in *FLVB* and PCR to screen for the insertion in *CYP55* (**Supplemental Fig. S3A**). We isolated three independent progenies exhibiting a *flvB* mutant-like chlorophyll fluorescence transient (*25*) that exhibited insertion of a paromomycin cassette in the $4^{th}$ intron of the *CYP55* gene (**Supplemental Fig. S3B**). The absence of the FLVB protein was then confirmed via immunodetection (**Supplemental Fig. S3C**). The three triple mutant progenies were named *flvB cyp55-1*, *-2*, and *-3*; CC-4533 was used as the reference strain for these mutants.

**Other algal strains and culture conditions.** All wild type strains of eukaryotic microalgae and the culture media used in this study are listed in **Supplemental Table S3**. Culture media recipes can be found on the website of the culture collection of algae from the Göttingen University (SAG):(https://www.uni-goettingen.de/en/culture+collection+of+algae+%28sag%29/184982.html). Strains of *C. reinhardtii, Galdieria sulfuraria and Chlorella variabilis* were cultivated at 25°C under continuous dim light (5-10 µmol photons $m^{-2}$ $s^{-1}$). Other strains were cultured at 25°C in day/night cycle (16 h/8 h) under low light (20 µmol photons $m^{-2}$ $s^{-1}$).

**Membrane Inlet Mass Spectrometry (MIMS) measurements.** Gas exchanges were monitored inside a water-jacketed and thermoregulated (25°C) measuring chamber (modified Hansatech $O_2$ electrode chamber) containing 1.5 mL of cell suspension as already described (*25*). Microalgal cells were harvested, centrifuged at 450 *g* for 3 min and resuspended in their growing medium. *C. reinhardtii* cells where resuspended at a final concentration of 100 µg chlorophyll $mL^{-1}$; for **Fig. 4** and **Supplemental Fig. S6.** Other microalgal species were resuspended at similar biomass contents. The mass spectrometer sequentially monitors gas abundances ($CO_2$, $N_2$, NO, $O_2$, $CO_2$ and $N_2O$) by automatically adjusting the magnet current to the corresponding mass peaks (m/z = 12; 28; 30; 32 and 44 respectively). Since both $CO_2$ and $N_2O$ have their maximal fragmentation peak at m/z=44, $CO_2$ amount was determined by its mass peak at m/z=12 and the contribution of $CO_2$ to the m/z=44 determined using standard fragmentation tables, thus allowing the determination of $N_2O$ amounts. Maximum gross $O_2$ production was measured as described in (*22*) using a saturating green light of 3,000 µmol photon $m^{-2}$ $s^{-1}$.

**DNA extractions and PCR amplification.** Total DNA was extracted using Chelex 100 (Sigma-Aldrich). Putative insertions were confirmed in the three putative *cyp55* mutant strains by PCR using *dreemTaq* DNA polymerase with GC Buffer (Thermo Scientific). 5 sets of primers (**Supplemental Table S2**) were designed according to Cre01.g007950 gene sequence (www.phytozome.jgi.doe.gov) to target the predicted insertion loci of the paromomycin resistance cassette (**Supplemental Fig. S1; Supplemental Table S2**). For the 13-f1/13-r1 primer set, PCR cycles were as follows: 2 min at 95°C / 40 cycles: 30 s at 95°C, 30 s at 60°C, 4 min 30 s at 72°C / 2 min at 72°C. For the two sets of primers 14-r2/14-f2 and 95-r2/95-f2, PCR cycles were as follows: 2 min at 95°C / 40 cycles: 30 s at 95°C, 30 s at 60°C, 2 min 30 s at 72°C / 2 min at 72°C. For the two sets of primers 14-r1/14-f1 and 95-

r1/95-f1, PCR cycles were as follows:2 min at 95°C / 40 cycles: 30 s at 95°C, 30 s at 58°C (14-r1/14-f1 ) or 60°C (95-r1/95-f1), 1 min at 72°C / 2 min at 72°C. PCR products were separated on 1.5% (w/v) agarose gels.

**Supplemental Table S1 | *cyp55* mutant strains from the Chlamydomonas Library Project (CLiP).** For each putative mutant strain, DNA sequences flanking the insertional cassette were downloaded from the CLiP website and blasted in Phytozome (www.phytozome.jgi.doe.gov) against *C. reinhardtii* genome (v5.5) to predict insertion sites. In *cyp55-13, cyp55-14* and *cyp55-95* strains, the flanking sequences were located only in the *CYP55* locus (Cre01.g007950).

| Strain accession number (CLiP) | Short name (this work) | Flanking sequence | Predicted locus |
|---|---|---|---|
| **LMJ.RY0402.126113** | *cyp55-13* | 1) TAATCGGCCCGCTCGGCAGACTTCCCCTTC<br>2) GAGAATGGGATATCAGCAACAGAAACTCAG | *CYP55* |
| **LMJ.RY0402.209714** | *cyp55-14* | 1) AAACACCCATGCACGTGTACCAATAAGTTC<br>2) ACCATCAGGCAGGGGAGTGGCAGGAAGTAA | *CYP55* |
| **LMJ.RY0402.177695** | *cyp55-95* | 1) CGTCATCTCACTGCTGCAGCACCCGGACCA<br>2) GCCCAGGTTGATCTGGGTGGCCACTGTGGC | *CYP55* |
| **CMJ030 (CC-4533)** | Control strain (WT) | | None |

**Supplemental Table S2 | Primers used to characterize *cyp55* mutants.** Primers were designed to target the different predicted insertion loci.

| Mutant alleles | Primer name | Primer sequence | Expected length of amplified DNA for a WT *CYP55* sequence (bp) |
|---|---|---|---|
| ***cyp55-13*** | 13-f1 | GGAGAAGCGGCCATCCTG | 349 |
| | 13-r1 | CCCACATGGACAAGGGTAAGC | |
| ***cyp55-14*** | 14-f1 | CACCTGCACGTCGGCCAC | 350 |
| | 14-r1 | CACTCCCGACCGTCCTATC | |
| | 14-f2 | GAACTTATTGGTACACGTGCATGG | 27 |
| | 14-r2 | CTGAACCTTACTTCCTGCCAC | |
| ***cyp55-95*** | 95-f1 | GTGGCAAGGGACGTTACG | 384 |
| | 95-r1 | GGCTGTGCGTGTAAGGTATG | |
| | 95-f2 | GTGCTGCAGCAGTGAGATG | 6 |
| | 95-r2 | GGCCACCCAGATCAACCTG | |

**Supplemental Table S3 | Eukaryotic algae used in this study and their respective genotype.** Strains were purchased from the following algal culture collections: Gottingen University (SAG), University of Texas (UTEX), Chlamydomonas Library Project (Clip), Culture collection of algae and protozoa (CCAP) or supplied by different laboratories. Shown are the origin and reference of each strain used and the GenBank accession number of the *CYP55*, *FLVA* or *FLVB* genes when present in the species genome. GenBank accession numbers are provided with a hypertext link to the reference sequence in NCBI (https://www.ncbi.nlm.nih.gov).

| Specie | Origin and reference number (if any) | CYP55 (if any) | Flvs (if any) | Culture medium | Genome reference |
|---|---|---|---|---|---|
| *Chlamydomonas reinhardtii (CC-4533)* | CLiP (CMJ030) | XM_001700220.1 | XM_001699293.1 XM_001692864.1 | TAP | (*54*) |
| *Chlorella variabilis NC64A* | Dr. Blanc (Marseille) | XM_005848783.1 | XM_005849680.1 XM_005849412.1 | TAP | (*55*) |
| *Coccomyxa subelispoidea* | SAG (216-13) | | XM_005643392.1 XM_005643394.1 | ES | (*56*) |
| *Tetraselmis subcordiformis* | SAG (161-1a) | | GBEZ01019221.1 GBEZ01009242.1 | SWES | (*57*) |
| *Galdieria sulfuraria* | UTEX (UTEX2919) | | | BG11 | (*58*) |
| *Porphyridium purpureum* | SAG (1380-1a) | | | Porph | (*59*) |
| *Nannochloropsis gaditana* | CCAP (CCMP527) | | | SWES | (*60*) |
| *Thalassiosira pseudonana* | Dr. Gontero (Marseille) | | | SWES +Si | (*61*) |
| *Phaeodactylum tricornutum* | Mr. Fleury (Cadarache) | | | SWES | (*62*) |

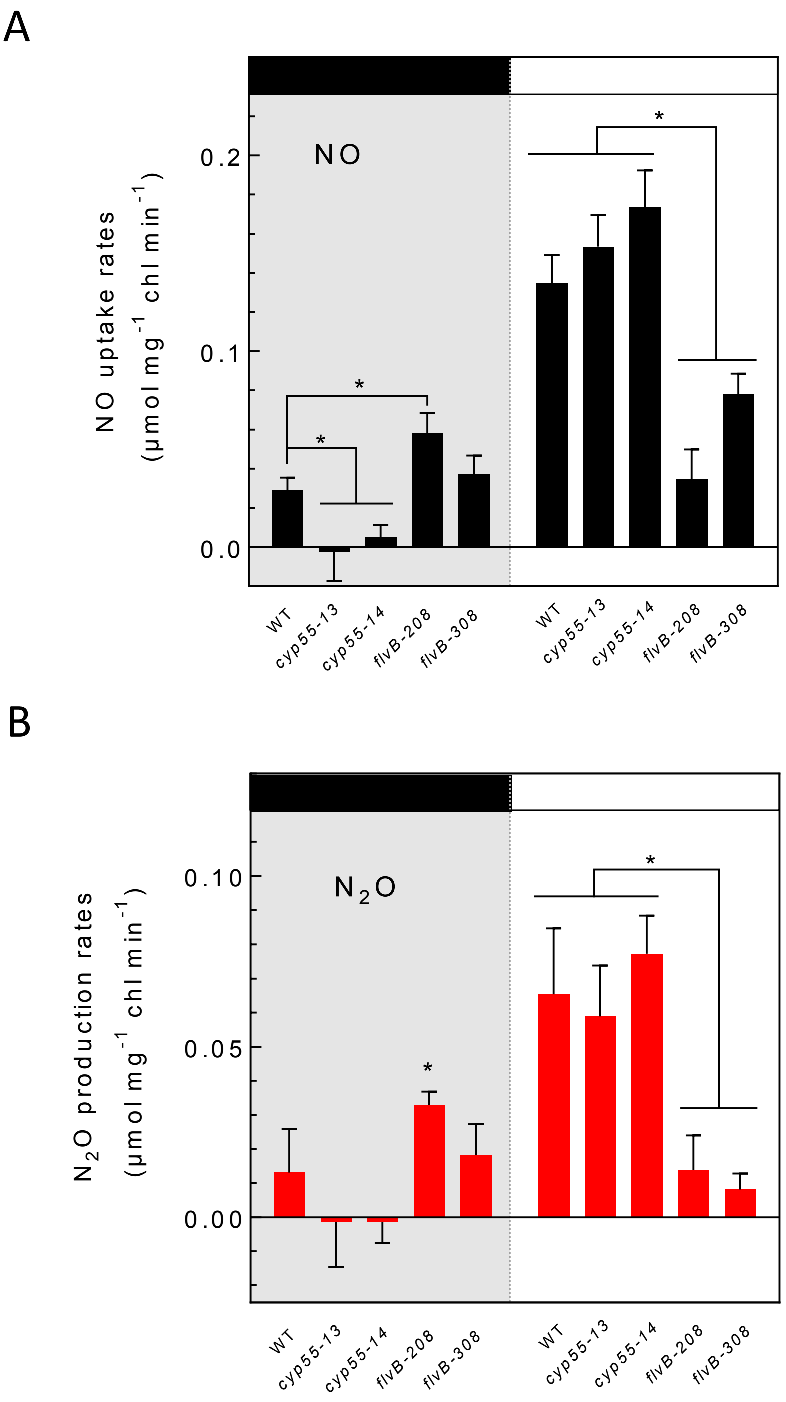


**Supplemental Figure S1. Photoreduction of NO into $N_2O$ in two additional *flvB* and *cyp55* mutants.** NO and $N_2O$ gas exchange were measured during a light transient as described in Fig. 1 in the control strain (WT), in two *cyp55* mutant (*cyp55-13* and *cyp55-14*) and in two *flvB* mutants (*flvB-208* and *flvB-308*). (A) dark (left panel) and light-dependent (right panel) NO uptake rates. (B) Dark (left panel) and light-dependent $N_2O$ production rates (right panel). Note that the *flvB-208* mutant showed a higher reduction of NO to $N_2O$ in the dark. This effect was not linked to the FLVB deficiency and could be due to an undetected random insertion of the paromomycin resistance cassette in the *flvB-208* genome (*27*). However, the light-dependent $N_2O$ production rate was diminished as observed in the two other *flvB* mutants. Data shown are mean values ± SD (n=4). Asterisks mark significant differences ($P < 0.05$) based on multiple T-tests.

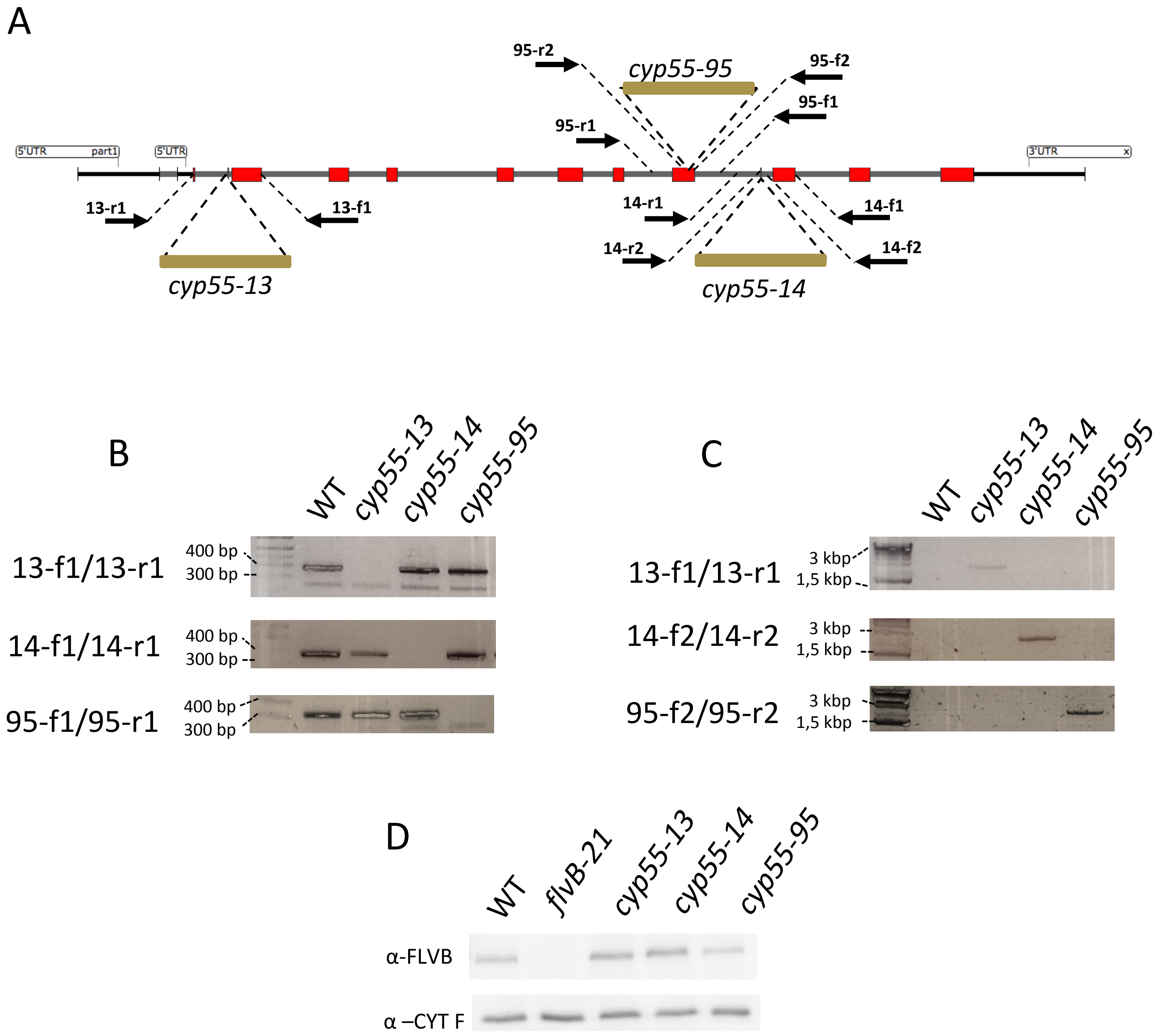


**Supplemental Figure S2. Characterization of *cyp55* mutants.** (A) Predicted insertion sites of the paromomycin resistance cassette in three putative *cyp55* mutants and location of primers used for PCR amplification. The length of the paromomycin resistance cassette is approximately 2,500 bp (*27*). (B) PCR verification of insertion sites of the three *cyp55* mutants using primers described in Supplemental Table S2. Primers were designed to amplify insertion loci of the resistance cassette. (C) PCR amplification of the insertion loci with the cassette. When amplification was impossible with primers used in (B), primers located closer to the predicted insertion loci were designed (Supplemental Table S2). (D) Immunodetection of FLVB and CYTOCHROME F (used as a loading control) in the WT strain, the *flvB-21* mutant, and the three *cyp55* mutants.

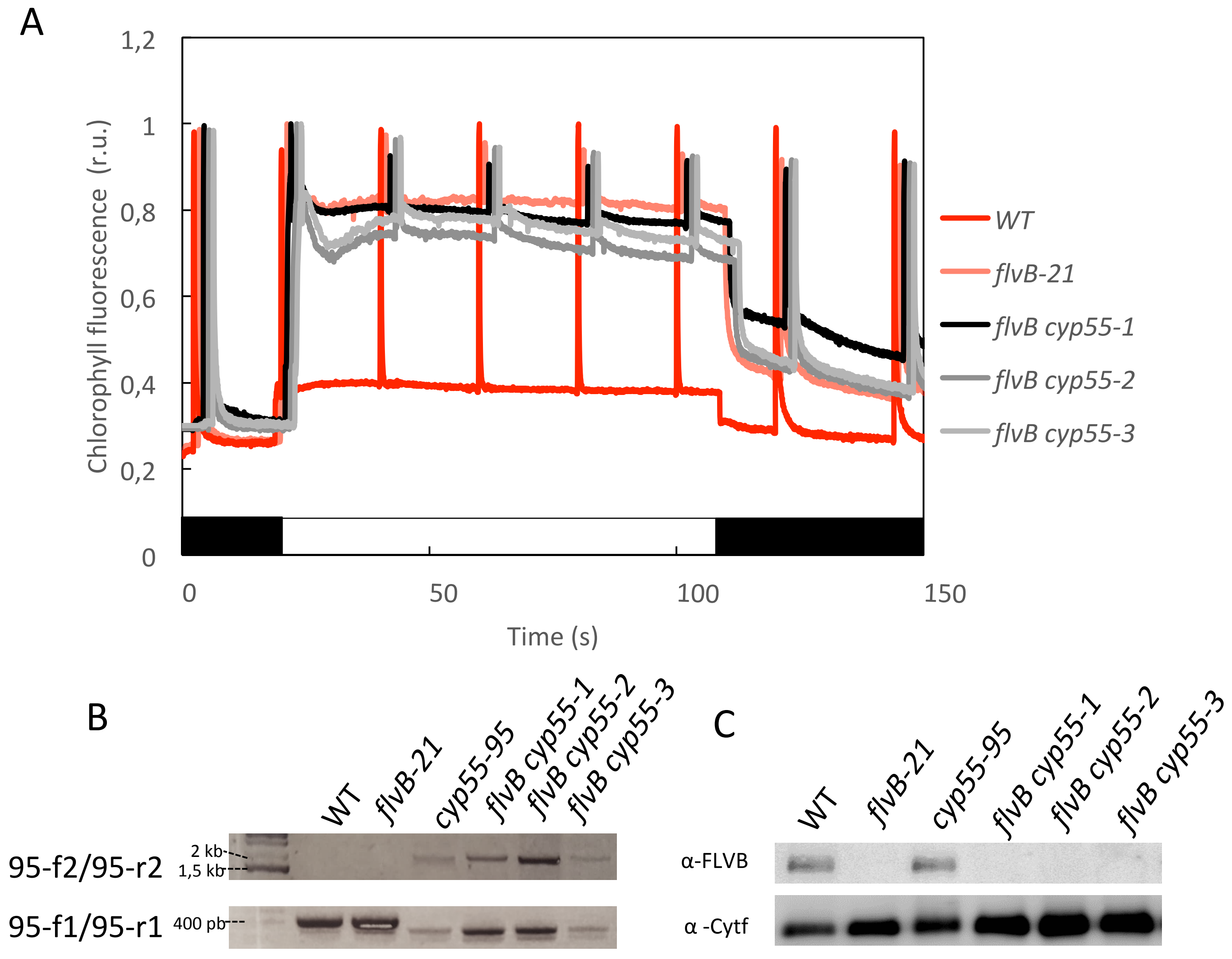


**Supplemental Figure S3. Characterization of *flvB cyp55* double mutants.** Upon crossing *flvB-21* and *cyp55-95* strains, progenies were screened for *FLVB* deficiency based on their chlorophyll fluorescence pattern as previously reported (*22, 25*), and then PCR analysed for an insertion in the *CYP55* locus. (A) Chlorophyll fluorescence pattern of a control strain (WT), *flvB-21* mutant and three independent progenies (*flvB cyp55-1, -2, -3*). Chlorophyll fluorescence measurements were performed using a pulsed amplitude modulated fluorimeter in the dark (black boxes) and under red actinic light (100 µmol photon $m^{-2}$ $s^{-1}$). Data are normalized to initial $F_M$ measurement and slightly shifted on the time axis for clarity. (B) PCR amplifications targeting the *cyp55-95* insertion locus. (C) Immunodetection of FLVB and CYTOCHROME F (used as a loading control) in the WT, parental *flvB-21* and *cyp55-95* strains, and in the three independent progenies (*flvB cyp55-1*, *-2, -3*).

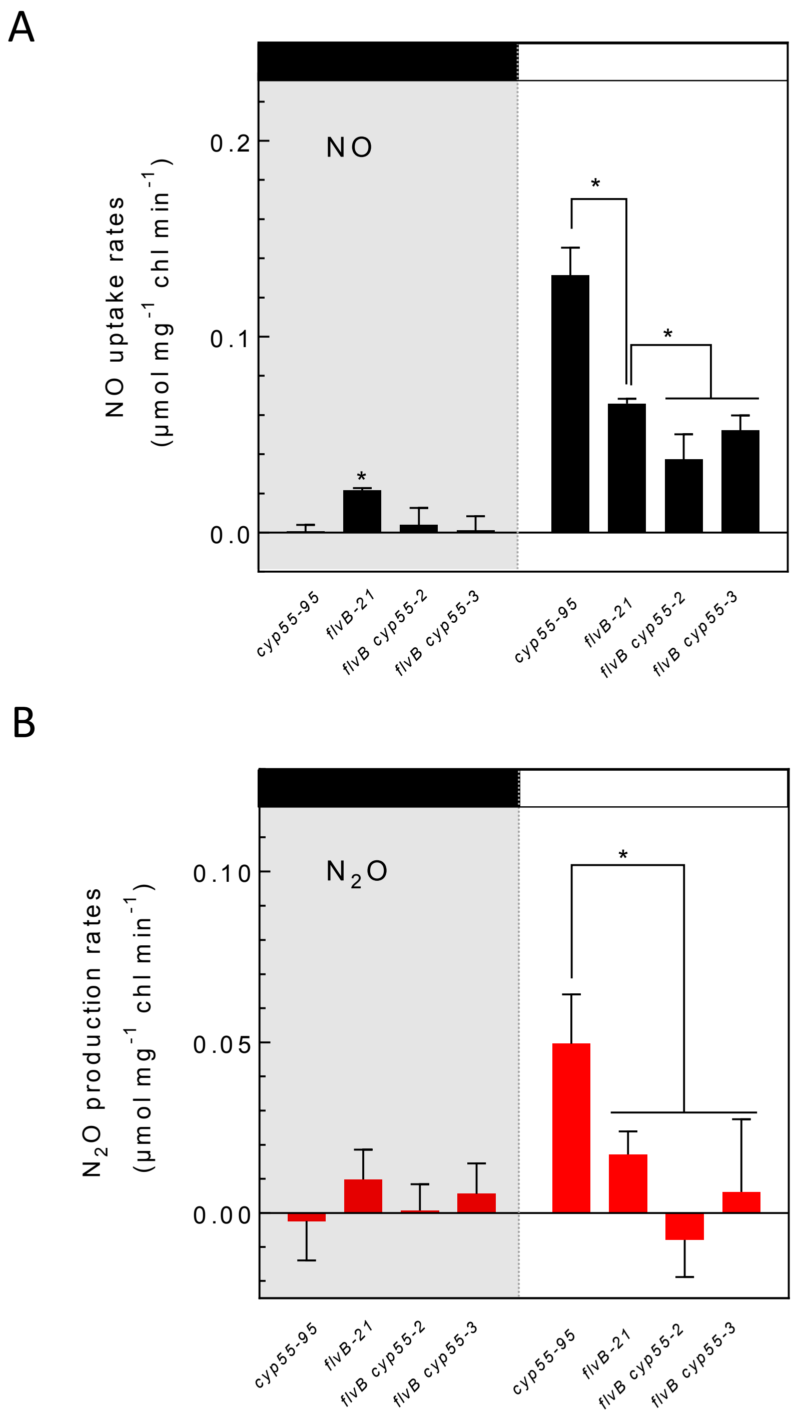


**Supplemental Figure S4. Photoreduction of NO into $N_2O$ in two additional *flvB cyp55* double mutants and their parental strains.** NO and $N_2O$ gas exchange were measured during a light transient as described in Fig. 1. (A) Dark (left panel) and light-dependent (right panel) NO uptake rates. (B) Dark (left panel) and light-dependent (right panel) $N_2O$ production rates. Data shown are mean values ± SD (n=4). Asterisks mark significant differences ($P < 0.05$) based on multiple T-tests.

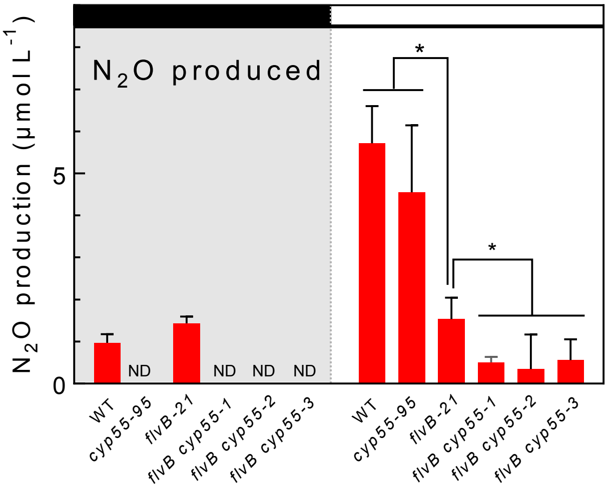


**Supplemental Figure S5. Integrated $N_2O$ amounts produced during a light transient.** NO and $N_2O$ gas exchange were measured as described in Fig. 1 in the control strain (WT), in simple mutants (*flvB-21* and *cyp55-95*) and double mutants (*flvB cyp55-1*, *-2*, *-3*). $N_2O$ production was integrated during one minute of darkness (left panel) and during the first minute of illumination. Light-dependent $N_2O$ production was calculated as the difference between light and dark productions (right panel). Data shown are mean values ± SD (n=4). Asterisks mark significant differences ($P <$ 0.05) based on multiple T-tests.

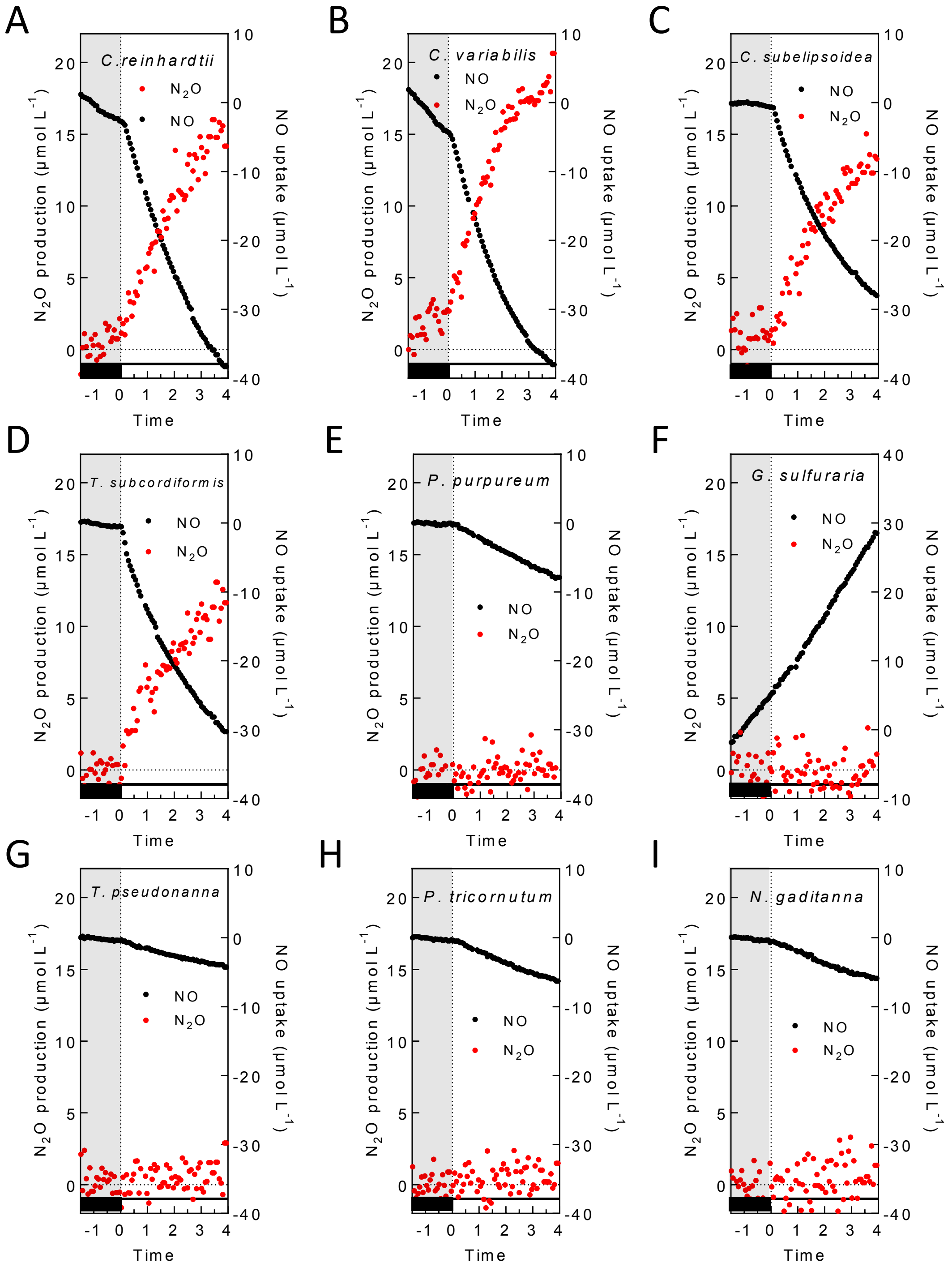


**Supplemental Figure S6. NO uptake and $N_2O$ production in various microalgal species during a dark to light transient.** NO and $N_2O$ gas exchange were measured during a light transient as described in Fig. 1 in cell suspensions of *Chlamydomonas reinhardtii* (A), *Chlorella variabilis* (B), *Coccomyxa subelipsoidea* (C), *Tetraselmis subcordiformis* (D), *Porphyridium purpureum* (E), *Galdieria sulfuraria* (F), *Thallassiosira pseudonana* (G), *Phaeodactylum tricornutum* (H) and *Nannochloropsis gaditana* (I). Shown are representative traces of cumulated amounts of NO uptake (black circles) and $N_2O$ production (red circles) measured in the different strains.
